# Mechanisms Driving Thoracic Aortic Aneurysm Stability

**DOI:** 10.1101/2025.10.20.683467

**Authors:** Erica L. Schwarz, David S. Li, Colin W. Means, Roland Assi, Jay D. Humphrey

## Abstract

Thoracic aortic aneurysms (TAAs) arise from a combination of biological and mechanical factors. Current clinical guidelines use size and rate of expansion to stratify risk, but such metrics do not predict if an aneurysm will stabilize, grow, dissect, or rupture. Computational biomechanical models can provide insights into mechanisms of aneurysm behavior that would be difficult or impossible to capture in vivo. Here, we use a constrained mixture theory of growth and remodeling to simulate lesion progression while co-varying rate-dependent parameters that contribute to the natural history of aneurysm growth. This includes insults to the material structure and mechanosensitivity of the vessel. This framework successfully simulates clinically-relevant phenotypes, including cases where lesions with initially similar degrees of dilatation or rates of expansion diverge in behavior later in their progression. By capturing this spectrum of outcomes, our framework lays a foundation for more accurate, patient-specific risk prediction and future integration of machine learning tools to accelerate translation into clinical practice.

## INTRODUCTION

Thoracic aortic aneurysms (TAAs) progressing to dissection and rupture are increasingly responsible for significant morbidity and mortality, with most of these lesions presenting at the aortic root or within the ascending aorta. Fortunately, advances in clinical screening of underlying pathogenic variants or congenital defects, as well as advances in prosthetic vascular graft technology and surgical technique, have improved treatment outcomes for patients harboring TAAs (Verhagen et al., 2018). Current clinical guidelines recommend prophylactic surgical replacement of the diseased segment of the thoracic aorta when either the maximal diameter exceeds a stated threshold value or the rate of enlargement exceeds a stated value, or both (Members et al., 2022). Nevertheless, many TAAs below recommended values dissect or rupture, while many larger lesions continue to be found incidentally without symptoms or tissue failure. There is, therefore, a need to understand better the natural history of TAAs and what underlying mechanisms determine their clinical destiny.

Computational biomechanical models promise to provide information beyond what is generally available clinically. Such models have advanced to account for many of the complexities of the thoracic aorta, including its complex geometry, nonlinear material properties, and hemodynamic loading. Notwithstanding the dominance of maximum diameter in risk assessment, there is increasingly more attention to rates of increase in lesion size (Henry et al., 2025; Oladokun et al., 2016). Among computational models that can account for rates of change of vascular geometry and material properties, constrained mixture models are particularly applicable. Briefly, the constrained mixture framework specifies three classes of constituent-specific constitutive relations (Humphrey and Rajagopal, 2002): one for the rate of mass production, one for the rate of mass removal, and one for the material properties of the existing constituents. The rates of production and removal depend on mechanobiological factors, including the ability of the cells to mechanosense their local environment, deviations between sensed and homeostatic values of mechanical metrics, and mechanosensitivity of the cellular response to these deviations.

The purpose of this work is two-fold. First, to quantitatively compare the computational utility of two different formulations of constrained mixture models — (i) a fully time-resolved approach versus (ii) a mechanobiologically equilibrated (MBE) approach — both of which can be coupled to hemodynamic solvers as needed, though with very different computational costs (Latorre and Humphrey, 2020a; Pfaller et al., 2024; Schwarz et al., 2023). Second, to contrast differential effects of different rate parameters that represent mechanobiological deficiencies and can dictate whether the growth of a TAA will arrest, continue slowly, or result in a mechanobiological instability indicative of functional or structural failure (Figure 1). As will be seen, the MBE approach allows efficient prediction of stable configurations in which an equilibrated state is possible, which provides a benchmark for an idealized, stable outcome. Through comparison of MBE predictions to fully time-resolved simulations, however, we show various mechanisms by which this stability can be lost, pointing to critical rate-dependent foci that influence the resulting geometry and mechanical vulnerability of the TAA.

**Figure 1:**
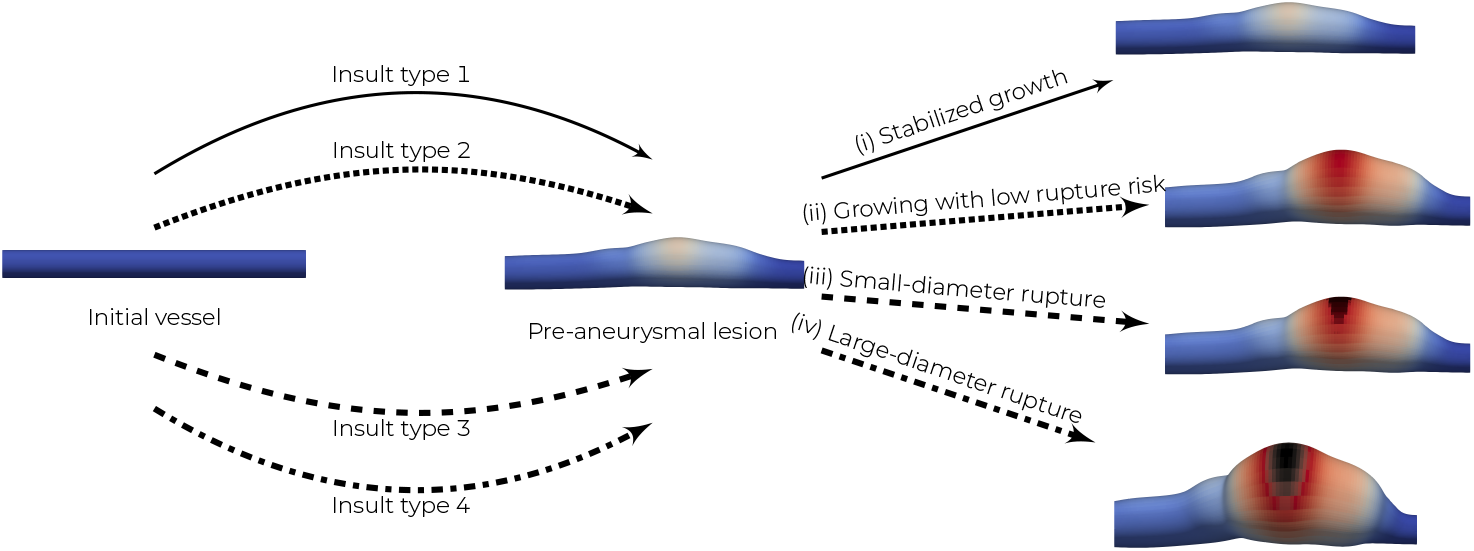
Illustration of possible outcomes of aneurysm progression. Different insult types contributing to disease progression can result in similar pre-aneurysmal lesions at intermediate times. Despite these initial geometrical similarities, continued progression can result in different outcomes, depending on the underlying mechanisms of insult. This includes: (i) stabilization of geometry and wall integrity, (ii) unstable growth with low risk for dissection/rupture, and growth with high risk for dissection/rupture either (iii) below or (iv) above stated clinical thresholds.

## METHODS

### Constrained mixture theory

Constrained mixture theory for soft tissues has been a widely adopted method for simulating aneurysm growth (changes in mass) and remodeling (changes in structure), centered on modeling tissue-level behavior by accounting for the production (deposition of matrix, proliferation of cells) and removal (degradation of matrix, death of cells) of individual vascular constituents. Full details of constrained mixture theory for vascular growth and remodeling (G&R) are described elsewhere (Humphrey, 2021). Briefly, constrained mixture theory posits that structurally significant vascular constituents possess individual natural configurations and have distinct material properties and turnover but are constrained to move with the mixture as a whole after deposition. Modeling the aortic wall as a tissue comprised of fibrillar collagens, smooth muscle cells, and elastin, we postulate that the apparent mass density *ρ*^*α*^ of individual constituents *α* at current time *s* can be expressed in terms of constitutive relations for mass production and removal (that is, survival) such that

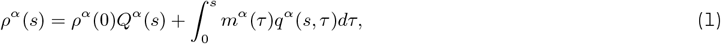

where *m*^*α*^(*τ* ) is the rate of mass density production at time *τ, q*^*α*^(*s, τ* ) is the fraction of constituent *α* deposited at time *τ* that survives until time *s*, and *Q*^*α*^(*s*) is the fraction of constituent *α* existing at time 0 that survives until time *s*. Previous studies have demonstrated the utility of stress-mediated relations for mass density production and removal (Humphrey, 2021). Traditionally, production *m*^*α*^(*τ* ) is modeled via a basal production value that can be modulated by changes in mixture-level stress, namely

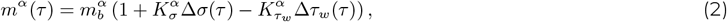

where 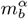 is a basal production rate and Δ*σ* and Δ*τ*_*w*_ represent the deviations in intramural and wall shear stress from homeostatic values, respectively, of the form

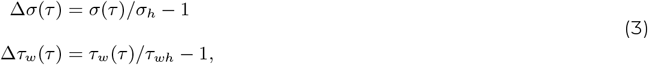

where *σ*(*τ*) is a scalar measure of the intramural stress at time *τ, τ*_*w*_(*τ* ) is the wall shear stress at time *τ, σ*_*h*_ is the homeostatic intramural stress, and *τ*_*wh*_ is the homeostatic wall shear stress, each denoted by *h*. 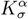 and 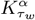 are gain-like parameters that stipulate the level of mechanosensitivity to the particular deviation in stress. Similarly, for mass removal,

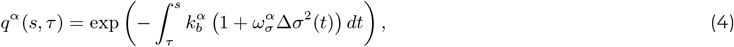

where 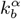 is a basal removal rate and 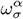 modulates the removal rate according to magnitude of the deviation in intramural stress. For simplicity, these gain-like parameters will be assumed to be equal for all the constituents but can be defined on a constituent-specific basis in the context of appropriate data.

The constituent strain energy can then be written as

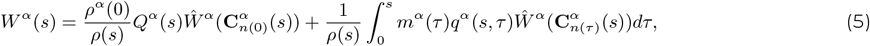

where 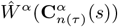 is a constituent-specific strain energy function and 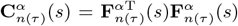 is the constituentspecific right Cauchy-Green tensor that arises from the constituent-specific deformation gradient 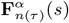, which represents the deformation of a constituent deposited at time *τ* at current time *s*. This is given as 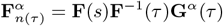, where **F**(*s*) is the deformation gradient of the mixture at *s*, and **G**^*α*^(*τ* ) is a tensor that accounts for pre-stretch of newly deposited constituents.

We then postulate that the mixture-level strain energy *W* (*s*) is given by a simple rule of mixtures such that, conceptually, *W* (*s*) =∑ _*α*_ *ϕ*^*α*^ (*s*)*W* ^*α*^ (*s*), where *ϕ*^*α*^ (*s*) = *ρ* ^*α*^ (*s*)*/ρ*(*s*) is the current constituent mass fraction at time *s, ρ*^*α*^ (*s*) is the apparent mass density of the constituent, and *ρ*(*s*) is the mass density of the mixture as a whole. From this we define the overall tissue-level Cauchy stress ***σ*** as

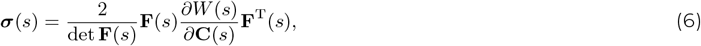

which allows efficient computation of the intramural stress in Equation 3, where 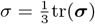.

We model the pseudoelastic mechanical behavior of collagen fibers (*α* = *c*) and passsive smooth muscle cells (*α* = *m*) as Fung-type exponentials with preferential fiber directions of the form

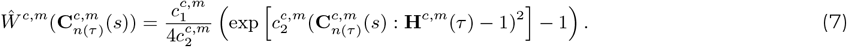

where 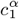 and 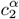 are material parameters and **H**^*c,m*^(*τ* ) = **h**^*c,m*^(*τ* ) *⊗* **h**^*c,m*^(*τ* ), where **h**^*c,m*^(*τ* ) is the orientation of the fiber at time *τ*, defined by 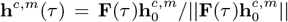, where 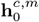 is the fiber direction in the reference configuration. This relationship specifies that new fibers are deposited preferentially in the direction of deformation. The deposition tensors are given by

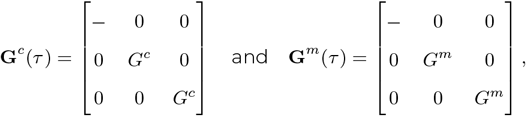

where *G*^*c*^ represents the pre-stretch of the collagen fiber at deposition, thought to be accomplished via integrin and actomyosin activity, and *G*^*m*^ is the pre-stretch of the smooth muscle cells (akin to cells spreading to reach a homeostatic mechanical state when placed on a substrate in vivo). Collagen fibers in the aortic wall naturally exhibit a splay of orientations ranging from circumferentially to axially oriented. This structural information can be captured by dividing collagen fibers into four subpopulations: circumferential (*θ*), axial (*z*), and symmetric diagonal (*d*), with associated fractions *{β*^*θ*^, *β*^*z*^, *β*^*d*^*}*. The diagonally oriented collagens exhibit an initial fiber angle *α*_0_ in the reference configuration relative to the axial direction, such that 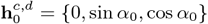. We also assume that collagen and smooth muscle cells have a maximum fiber age *A*_*max*_ during their turnover that captures the diminishing structural contribution of fibers and cells after significant removal. Effectively, this means that mass and material behavior from constituents that turn over (e.g. collagen, smooth muscle cells) is only considered, and subsequently integrated in Equation 1 and Equation 5, if deposited on the time interval [*s − A*_*max*_, *s*]. Lastly, we model elastin (*α* = *e*) as a neo-Hookean material of the form

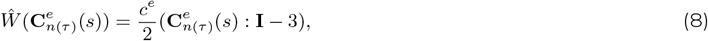

where *c*^*e*^ is a material parameter and **G**^*e*^(*τ* ) a three-dimensional deposition stretch given by

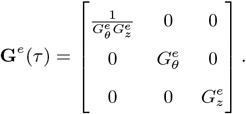

We note, given the considerably long half-life of elastic fibers in the aorta (on the order of decades), that elastin is assumed to not undergo natural turnover in the modeling framework. However, as discussed below, its properties can be compromised throughout the progression of TAA.

Linearization of the constrained mixture formulation consistent with these constraints enables efficient implementation within a finite element framework, where simultaneous solution of mechanical and mechanobiological equilibrium can be ensured across load steps that capture evolving geometries, compositions, and properties of interest for complex boundary value problems (see Schwarz et al. (2023) for detail on the full constrained mixture theory within a finite element framework). For this study, this methodology was implemented in the open-source FEBio solver. The source code can be found on GitHub (Schwarz, 2025).

### Mechanobiologically equilibrated formulation of constrained mixture theory

Constrained mixture theory for vascular G&R is motivated by the concept of mechanical homeostasis (Humphrey et al., 2015), which aims for balanced constituent turnover to achieve an unchanging mechanical state in which cells deposit new matrix with a preferred pre-stress. We have shown that this constrained mixture framework can be simplified via the assumption of mechanobiological equilibrium at long time scales (Latorre and Humphrey, 2018a,b), with homeostatic (steady-state) production balancing removal. Since homeostasis requires that the mixture mass, composition, and properties are unchanging, we observe from Equation 2 that 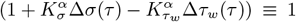 in mechanobiologically equilibrated configurations. In other words, 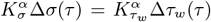. Additionally, we define the homeostatic deformation gradient **F**_*h*_ describing the deformation between the initial state and the long-term homeostatic configuration, along with its Jacobian *J*_*h*_ = det **F**_*h*_. It follows that, for mixture constituents that are assumed to not turn over (e.g., elastin), because their mass remains constant, the homeostatic mass fraction is given by

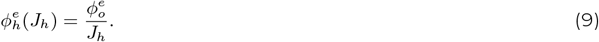

For constituents that are allowed to turn over (e.g., smooth muscle cells and collagen fibers), their evolved mass fractions can be computed relative to each other using

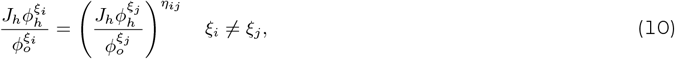

where *η*_*ij*_ is a turnover ratio between constituents *ξ*_*i*_ and *ξ*_*j*_ . We note that, due a lack of experimental observations to determine this value in disease cases, we let *η*_*ij*_ = 1 for this study. Similarly, since the intramural and wall shear stress contributions to mass production balance each other, one need only specify the ratio between the gain parameters 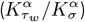 as an input, as done previously (Latorre and Humphrey, 2020b). This framework has also been implemented in FEBio as a custom material plugin (Latorre and Humphrey, 2020a,b) and provides rapid, path-independent estimation of the long-term evolved state of the full constrained mixture model with a simulation time on the order of seconds. Importantly, this also serves as a benchmark for a fully stabilized TAA progression, facilitating key comparisons in rate-dependent mechanisms discussed below.

### Mechanobiological insults contributing to aneurysm progression

To model how combinations of material integrity and mechanobiological sensitivity impact aneurysm progression and stability, we model TAAs by prescribing a time-varying mechanobiological insult *ϑ ∈* [0, 1] to an initially non-dilated vessel that is pre-loaded to in vivo conditions. The insult profile allows specific control of localized properties that induce deviations in mechanical and mechanobiological equilibrium to initiate aneurysmal dilatation and is defined using a modified expression from Latorre and Humphrey (2020b):

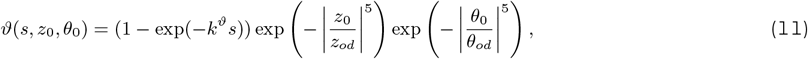

where *s* is the current G&R time, *z*_0_ and *θ*_0_ are axial and circumferential coordinates in the initial configuration, respectively, *k*^*ϑ*^ is a rate parameter representing the rate of insult progression over time, and *z*_*od*_ and *θ*_*od*_ represent the spread of the insult in the axial and circumferential directions, respectively, understanding that the maximum insult is centered on *{z*_0_, *θ*_0_*}* = *{*0.0, 0.0*}*.

Contributors to TAAs are thought to include disrupted material structure, altered material turnover rates, and compromised sensitivity to changes in hemodynamic environment. We focus here on aneurysms associated with losses in elastic fiber integrity, whereby the elastin modulus *c*^*e*^ in Equation 8 is reduced by a specified percentage, as done previously to model aneurysm formation (Goswami et al., 2022; Latorre and Humphrey, 2020b). Extending this approach, we further hypothesize that the sensitivity of vascular cells to changes in their local mechanical environment via integrin and actomyosin activity can be compromised in TAAs over time, leading to aberrant constituent production and removal. We thus use Equation 11 to parameterize gradual localized insults to elastic fiber integrity (*ϑ*^*e*^) and mechanosensitivity to intramural stress (*ϑ*^*p*^ for intramural stress-mediated production and *ϑ*^*r*^ for removal) and wall shear stress (*ϑ*^*τw*^ ) by assigning the normalized insult profile to affect multiple contributors at once. Specifically,

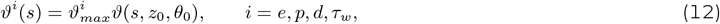

where the maximal severity value 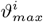 for each contributor is reached as *s → ∞. k*^*ϑ*^ in Equation 11 can then be tuned such that the insult severity reaches 99% of its maximum value at any point within the simulation time-course. Specifically, *ϑ*^*e*^ describes the percent reduction in the elastin modulus from its baseline value, that is, *c*^*e*^(*τ* ) = (1 *− ϑ*^*e*^(*τ* )) *c*^*e*^(0). Compromised mechanosensitivity of constituent production and removal are achieved by modifying the gain-like parameters in Equations 2 and 4 as

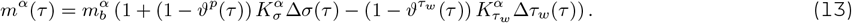

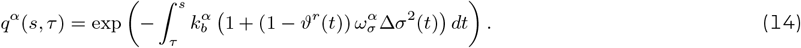

We assume the combined effects change together both temporally and spatially, governed by *k*^*ϑ*^, such that 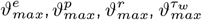 are reached at the same simulation time-step.

We model the aorta using material parameters (Table 1) based on biaxial biomechanical data of a normal thoracic aorta from a male wild-type mouse (Latorre and Humphrey, 2020b). Using the above definitions and parameters as inputs, the finite element solver was used to compute the displacement field for each aortic geometry associated with the mixture-level stress in the context of applied boundary conditions. The aortic domain was modeled as a uniform thick-walled cylinder and meshed using 27-node, fully-integrated quadratic hexahedral elements, with 1 *×* 160 *×* 40 elements along the radial, circumferential, and axial directions, respectively. Aortas were simulated at a fixed pressure of 105 mmHg, corresponding to an elevated in vivo mean arterial pressure. Where appropriate, we used symmetry conditions to calculate full-domain configurations from domain subsections. We compared outcomes of synthetic TAAs generated using full and equilibrated constrained mixture theory, noting that the equilibrated formulation determines the final configuration of the aneurysm directly and therefore only considers the final (maximum) value 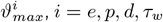 (Figure 2).

**Table 1:** Simulation parameters for aortic material and turnover properties. Initial values are for a normal wild-type male mouse descending thoracic aorta, derived from previous studies (Goswami et al., 2022; Latorre and Humphrey, 2018b, 2020b; Wilson et al., 2013). SMC = smooth muscle cells.

| **Material and Turnover Properties** |  |  |
| --- | --- | --- |
| Initial radius, thickness, length (mm, mm, mm) | $a_0, h_0, l_0$ | 0.6468, 0.042, 15.0 |
| Initial volume fractions (–, –, –) | $\phi_0^e, \phi_0^m, \phi_0^c$ | 0.34, 0.33, 0.33 |
| Directional collagen fractions (–, –, –) | $\beta^\theta, \beta^z, \beta^d$ | 0.056, 0.067, 0.877 |
| Initial diagonal collagen orientation (°) | $\alpha_0$ | $\pm 29.9$ |
| Elastin material property (kPa) | $c^e$ | 89.71 |
| Elastin deposition stretches (–, –, –) | $G_r^e, G_\theta^e, G_z^e$ | $1/(G_\theta^e G_z^e), 1.90, 1.62$ |
| Collagen material properties (kPa, –) | $c_1^c, c_2^c$ | 234.9, 4.08 |
| Collagen deposition stretch (–) | $G^c$ | 1.25 |
| Collagen degradation rate (1/day) | $k_b^c$ | 0.142 |
| Collagen production gains (–, –) | $K_\sigma^c, K_{\tau_w}^c$ | 1.0, 0 |
| Collagen degradation gain (–) | $\omega_\sigma^c$ | 1.0 |
| SMC material properties (kPa, –) | $c_1^m, c_2^m$ | 261.4, 0.24 |
| SMC deposition stretch (–) | $G^m$ | 1.20 |
| SMC death rate (1/day) | $k_b^m$ | 0.142 |
| SMC production gains (–, –) | $K_\sigma^m, K_{\tau_w}^m$ | 1.0, 0 |
| SMC death gain (–) | $\omega_\sigma^m$ | 1.0 |
| Maximum fiber age (days) | $A_{max}$ | 64 |

**Figure 2:**
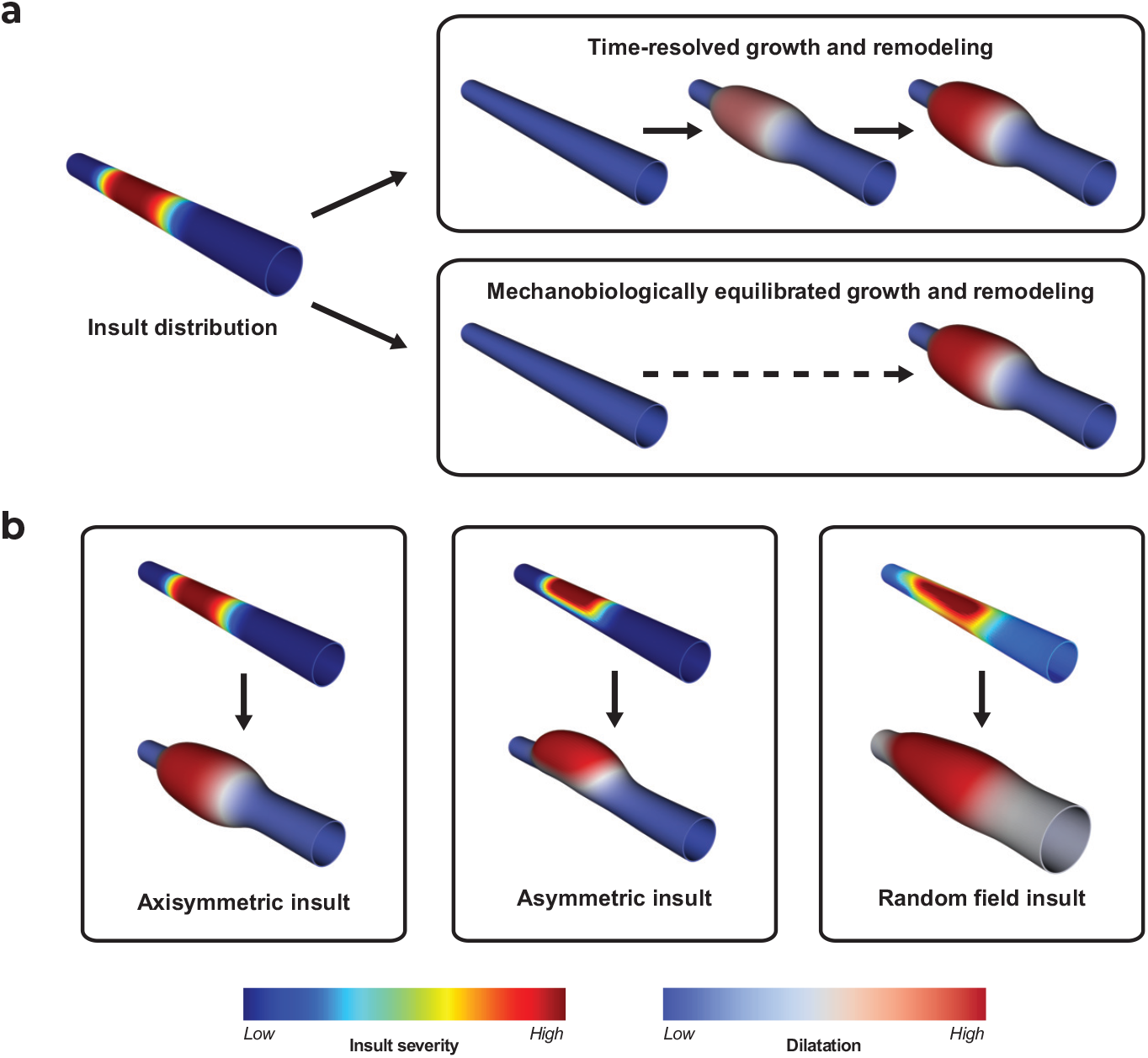
TAA simulation pipeline and mechanobiological insult profiles. (a) TAAs initiated by localized mechanobiological insults, defined by two-dimensional insult profiles, can be simulated using time-resolved or mechanobiologically equilibrated (MBE) finite element frameworks, which can be compared. Time-resolved simulations produce a time course of lesion growth, whereas MBE simulations identify a single stabilized long-term configuration, if such a configuration exists. (b) Illustrative examples of spatially axisymmetric, asymmetric, and random insult profiles. Shown are the prescribed spatial distribution (above, colored by severity) and the corresponding final TAA geometry (below, colored by dilatation).

### Spatial distribution of insult

Using fully time-resolved and mechanobiologically equilibrated constrained mixture formulations, we generate TAAs resulting from combinations of lost elastic fiber integrity and compromised mechanosensitivity for a wide range of insult profile types. To understand relationships among the spatial distribution of insults, their severity, and their rate dependence, we focus here on three classes of insult profiles (Figure 2).

Axisymmetric profiles vary only axially, assuming full circumferential spread and removing dependence on *θ*, such that Equation 11 simplifies to 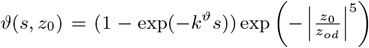. These are generated with a fixed axial spread *z*_*od*_ = 3.0 mm and maximum value (denoted the insult apex) centered at *z*_0_ = 0.0 mm.

Asymmetric profiles exhibit variation in both axial and circumferential directions as described by Equation 11. The circumferential spread is set to *θ*_*od*_ = *π/*2 with the apex at *θ*_0_ = 0, which limits the spread of the insult to only a portion of the full circumference in addition to the same axial spread as axisymmetric profiles.

Random insult profiles are generated by superimposing multiple (*N* = 4) asymmetric sub-profiles in the aortic domain. We define each sub-profile *f*_*i*_(*z*_0_, *θ*_0_) as

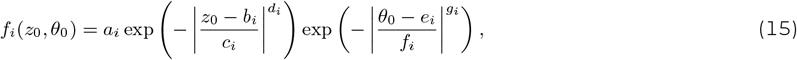

where *{a*_*i*_, *b*_*i*_, *c*_*i*_, *d*_*i*_, *e*_*i*_, *f*_*i*_, *g*_*i*_*}* are shape parameters independently sampled from a truncated normal distribution 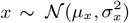 subject to *L*_*x*_ *≤ x ≤ U*_*x*_, where *{L*_*x*_, *U*_*x*_*}* denote lower and upper truncation bounds, respectively. Importantly, this method enables repositioning of the maximum insult within the aortic domain (*b*_*i*_ and *e*_*i*_) as well as variation in the spread (*c*_*i*_ and *f*_*i*_) and boundary softness (*d*_*i*_ and *g*_*i*_)of the insult region, all on a per-term basis. To ensure admissible profiles, the parameter bounds are chosen to produce similarly sized insult regions as the axisymmetric and asymmetric cases (i.e., constrained to be generally positive, with axial and circumferential spread distribution means centered on 5 mm and *π*, respectively.

The final random insult profile *ϑ*^*rand*^(*s, z*_0_, *θ*_0_) is computed by combining the terms as

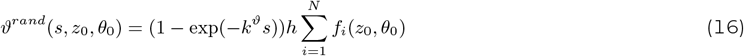

and used in place of Equation 11 in the simulations. *h* is a final normalizing parameter that imposes the maximum value of *ϑ*^*rand*^(*s, z*_0_, *θ*_0_) in the vessel domain is 1 as *s → ∞* (Figure S1). The parameters used in to generate insult profiles are summarized in Table 2.

**Table 2:**
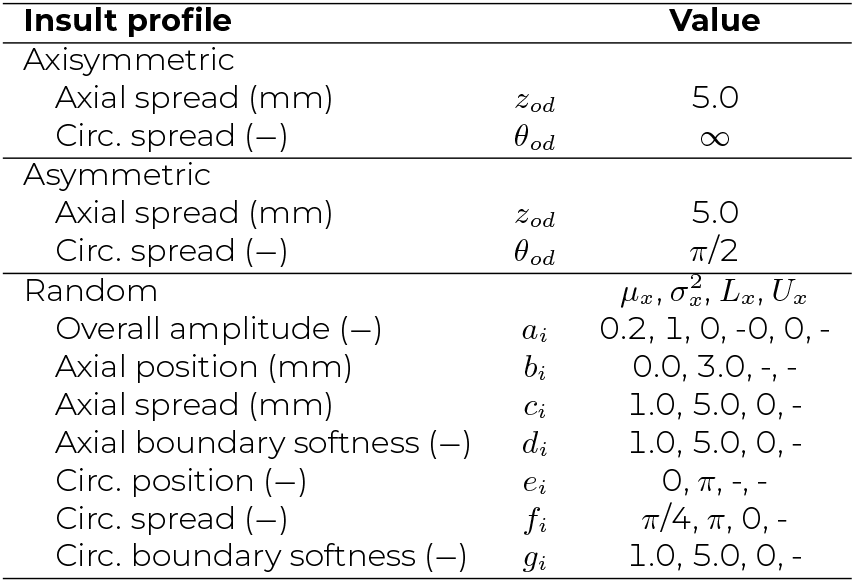
Spatial parameters of insult profiles. Summary of the shape parameters that define the insult profiles serving as input into the finite element simulations. For randomly generated insult profiles, the means (*µ*_*x*_), standard deviations 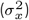, lower bounds (*L*_*x*_), and upper bounds (*U*_*x*_), of the truncated normal distribution are listed for each parameter.

### Numerical experiments

For illustrative purposes, and because of endothelial dysfunction in most TAAs, we focus on exploring the impact of insults to the mechanosensitivity of intramural stress deviations only (that is, 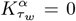 in Equation 13 and *ϑ*^*τw*^ = 0 for the time-resolved constrained mixture model, and 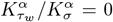 for the equilibrated), which allows direct comparison of the time-resolved and MBE results for a given set of input parameters. We performed experiments by varying pairs of insult severity parameters while holding the others constant for each class of insult profile:

**a** Elastic fiber integrity and production mechanosensitivity: co-varying 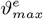 and 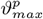 while fixing 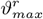 and *k*^*ϑ*^.

**b** Production and removal mechanosensitivity: co-varying 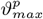 and 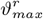 while fixing 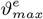 and 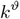.

**c** Elastic fiber integrity and insult rate: co-varying 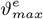 and *k*^*ϑ*^ while fixing 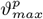 and 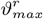.

For the time-resolved approach, we simulate a 720-day time-course (corresponding to the roughly 2-year lifespan of a mouse) to allow the time-resolved insults to equilibrate, if applicable, for comparison with the MBE. Quantities of interest (inner radius, wall thickness, circumferential material stiffness and stress, strain energy, and (scalar) intramural stress, all indications of aortic strength and function) are reported at the location of maximum insult amplitude, corresponding to the location of maximum diameter. Comparisons of progressive luminal expansion and wall thickening are particularly useful given their impact on wall stress as well as growth stabilization (Baek et al., 2005). We varied *k*^*ϑ*^ from [0.010, 0.050] and chose maximum insult values 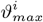 separately depending on the symmetry of the insult profile in order to identify combinations at which TAA stability would be lost.

## RESULTS

All insult profiles resulted in fusiform dilatations in the affected region of the model domain. We categorized time-resolved simulation outcomes according to the following four criteria:

- Stabilization: final growth rate at the apex *<*0.1 mm/year.
- Growing: final growth rate at the apex *≥*0.1 mm/year.
- Small-diameter rupture: the simulation exits prior to the full 720-day time-course and the final maximum radius is *<*1.5 times the original radius.
- Large-diameter rupture: the simulation exits prior to the full 720-day time-course and the final maximum radius is *≥*1.5 times the original radius.

In the context of this framework, the simulation exited when constituent removal 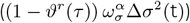 significantly exceeded production 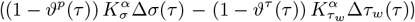, which were computed from mechanosensitivity parameters, mechanosensitivity insults, and intramural stress (Figures S2, S3, and S4). In a biological context, this exit corresponds to a catastrophic positive feedback loop whereby removal (e.g., degradation of matrix, death of cells) drastically increases, resulting in unbounded thinning of the wall, rapid increases in stress and stiffness, and expected rupture. While this created numerical non-convergence in the simulation framework, it is also directly analogous to biomechanical instability, making it an ideal metric to indicate aneurysm rupture.

Visualizing outcomes of the co-varying parametric simulations in a classification map facilitates identification of locations of multiple stability regimes within the parameter spaces (Figure 3 with these results evident from results described below for Figure 4, Figure 5, Figure 6, Figure 7, and Figure 8). This provides a framework for relating theoretical mechanobiological parameters to clinical quantities of interest and patient outcomes. In addition, the time-resolved approach provides critical knowledge on intermediate states of unstable TAAs, which are discussed below, focusing first on axisymmetric insult profiles.

**Figure 3:**
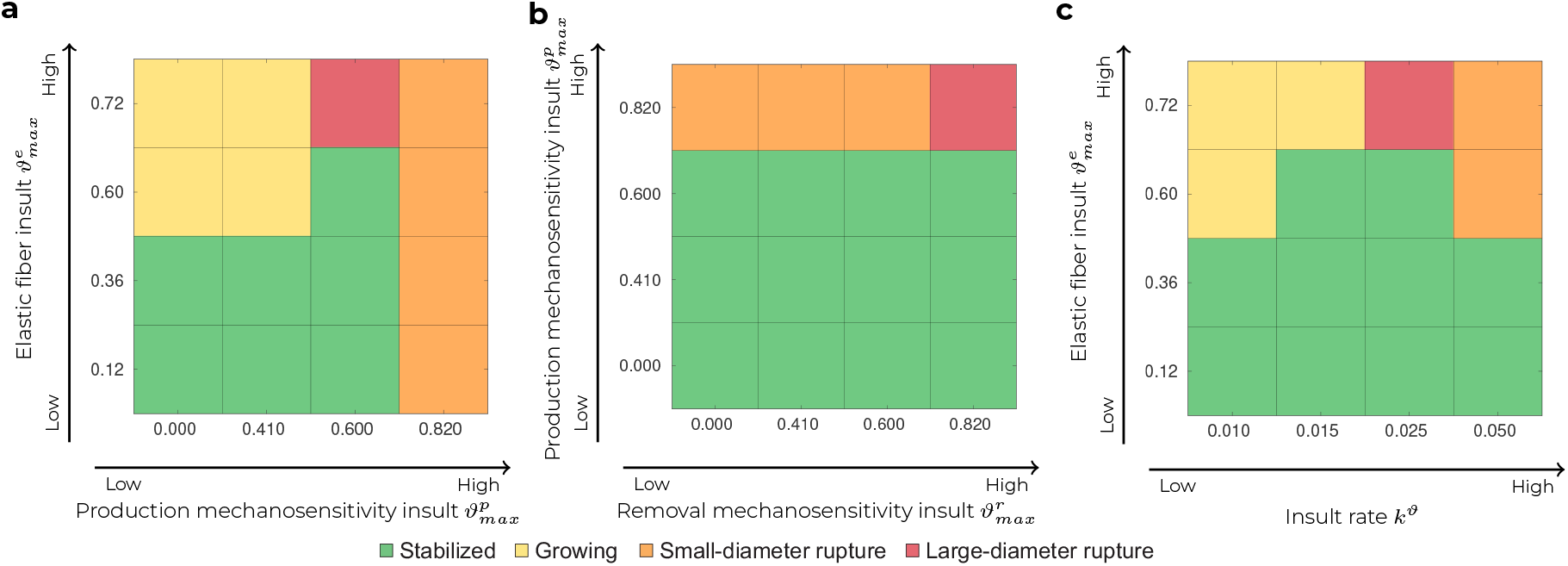
Co-varying biomechanical insults produce varying regimes of stability, growth, and rupture. Mechanobiological insult parameters 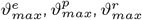, and *k*^*ϑ*^ are varied in pairs, and simulation outcomes are categorized according to the final lesion size and growth rate for converged simulations, or whether the simulation reached the rupture exit threshold. Green regions represent stabilized vascular outcomes whose configurations were simulated equally well using the MBE framework. (a) 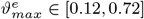 versus 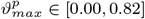 with 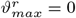 and *k*^*ϑ*^ = 0.025. Increased elastic fiber insult leads to unbounded growth, while increased production mechanosensitivity insult leads to rupture. (b) 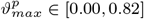 versus 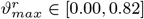 with 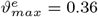 and *k*^*ϑ*^ = 0.05. Rupture potential is primarily driven by production rather than removal mechanosensitivity insult. (c) 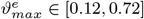 versus *k*^*ϑ*^ *∈* [0.010, 0.050] with 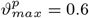 and 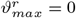. Higher insult rate increases rupture potential. Results are shown for an axisymmetric insult distribution.

**Figure 4:**
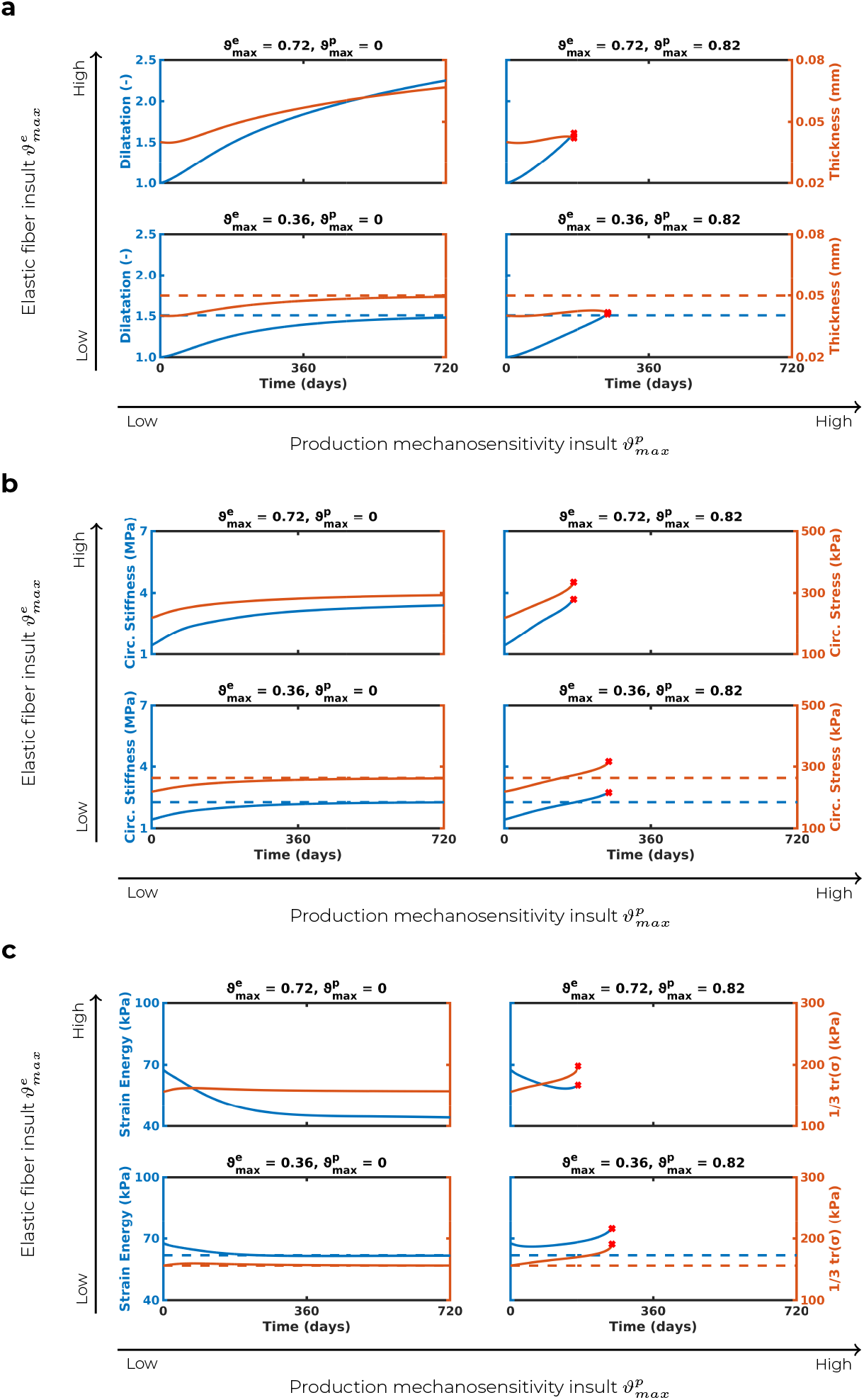
Effects of co-varying elastic fiber insult 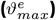 and production mechanosensitivity insult 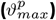. (a) Dilatation and thickness, (b) circumferential stiffness and stress, and (c) strain energy and intramural stress are shown over time for the fully time-resolved (solid lines) and MBE (dashed lines) models, evaluated at the location of maximum insult severity. Removal mechanosensitivity insult is fixed at 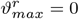, and insult rate is fixed at *k*^*ϑ*^ = 0.010. Increasing 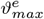 with low 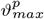 eventually results in an unstable growing aneurysm in which stiffness and stress no longer approach steady-state values but rather increase indefinitely. Increased 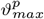 can destabilize a moderate 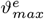 that is otherwise stable when 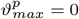. This results in increasing stiffness and stress over time that triggers the rupture exit threshold (red crosses). Both moderate and severe *ϑ*^*e*^ reach this rupture exit threshold, though they demonstrate strikingly different trends in strain energy at the exit point. Results are shown for an axisymmetric insult distribution.

**Figure 5:**
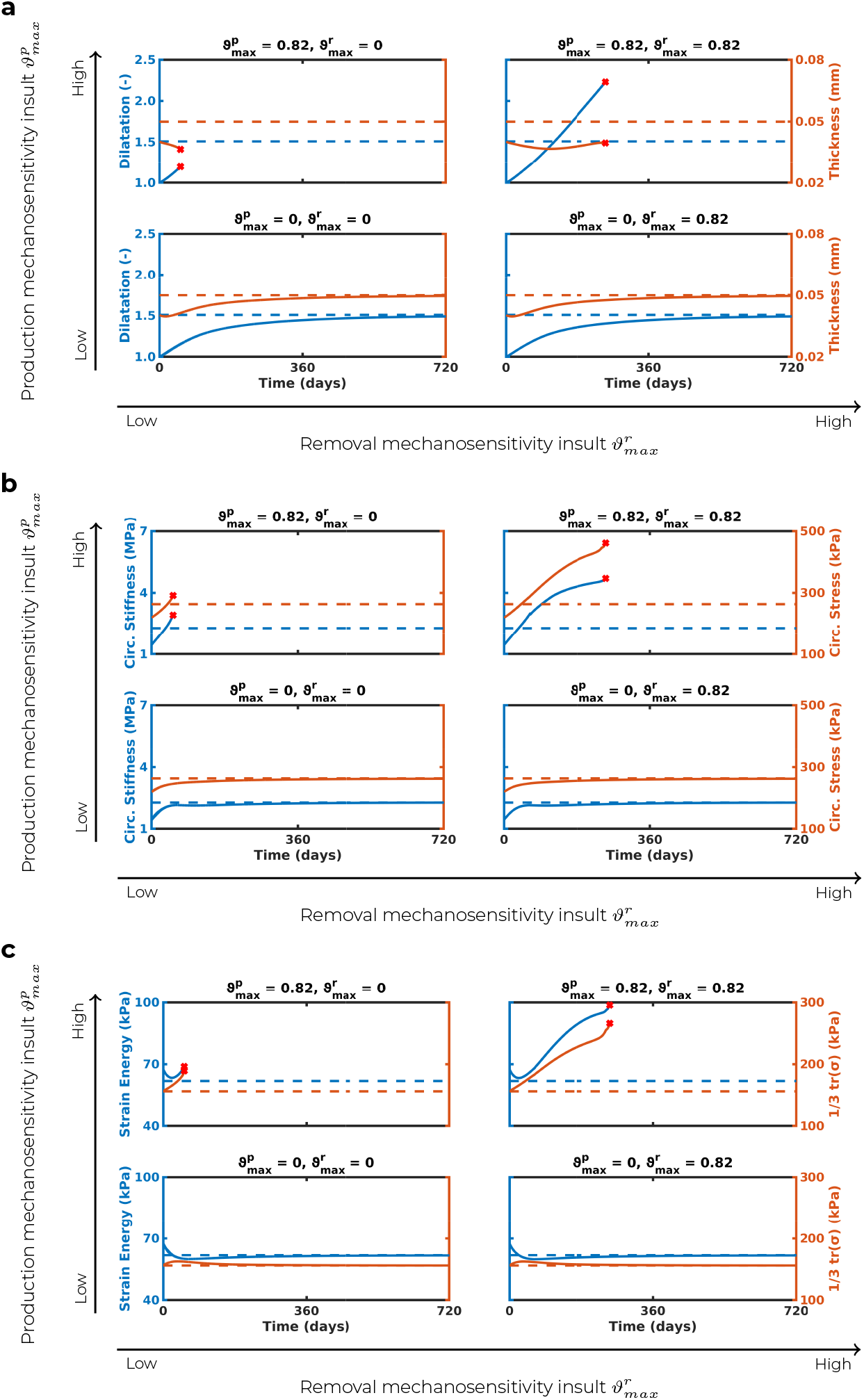
Effects of co-varying production 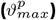 and removal mechanosensitivity insult 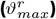. Similar to Figure 4 but with elastic fiber insult fixed at 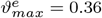 and insult rate fixed at *k*^*ϑ*^ = 0.05. TAA stability is maintained with low 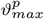 despite increases in 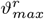. Conversely, high 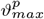 results in eventual loss of stability regardless of 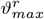 value, although increasing 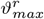 can delay the time of reaching the rupture exit threshold (red crosses). In addition to this delay in rupture, high–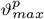 and 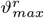 aneurysms exhibited significantly higher dilatation, circumferential stress and stiffness, and strain energy at rupture than their low–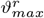 counterparts. Results are shown for an axisymmetric insult distribution.

**Figure 6:**
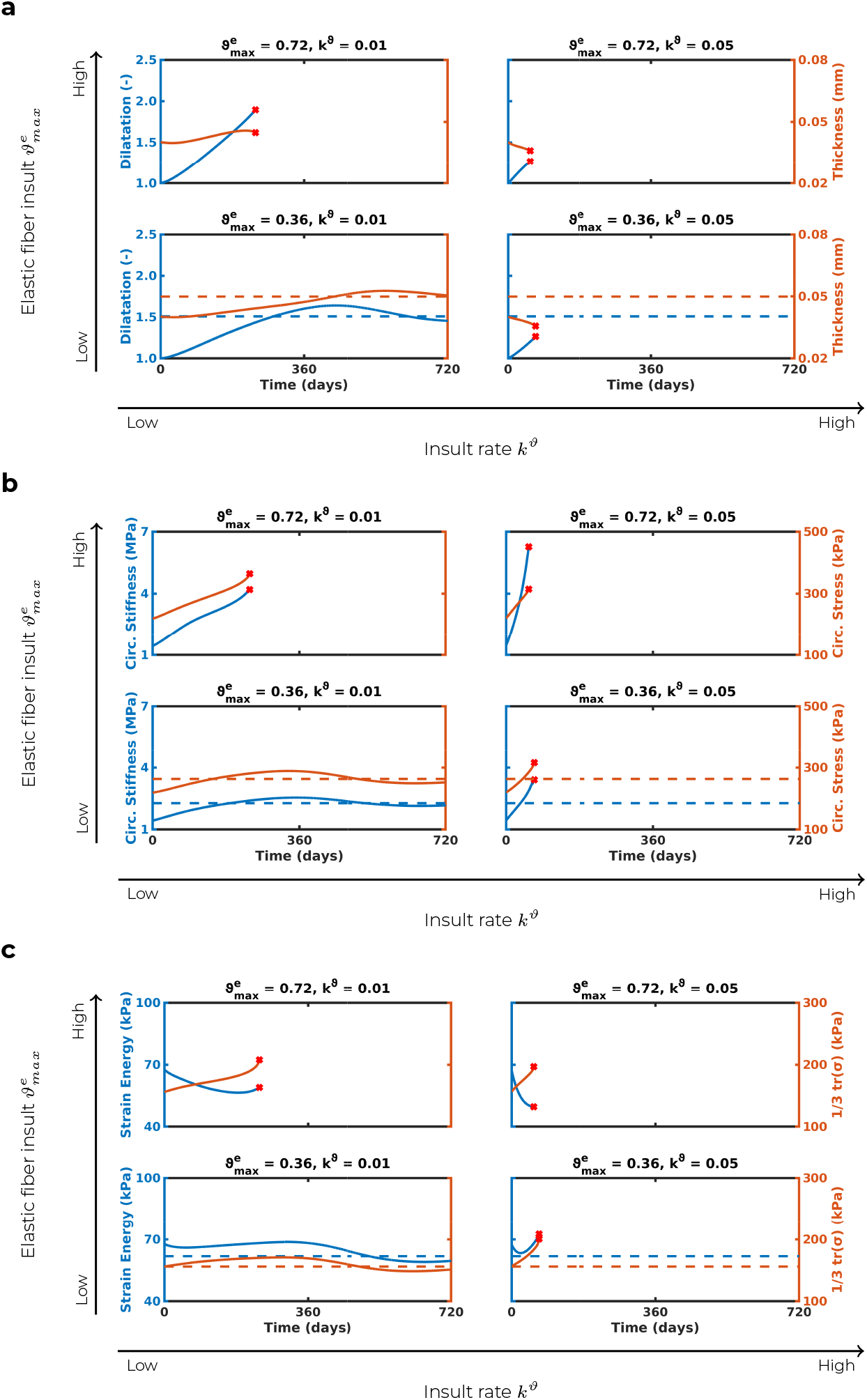
Effects of co-varying elastic fiber insult 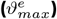 and insult rate 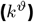. Similar to Figure 4 but with production mechanosensitivity insult fixed at 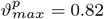 and removal mechanosensitivity insult fixed at 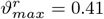. A rapid insult rate (high *k*^*ϑ*^) results in loss of convergence in which stiffness and stress rapidly increase beyond the MBE predictions, even at a moderate value of 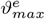 that would be expected to stabilize at low *k*^*ϑ*^ values. At severe 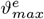 values, low values of *k*^*ϑ*^ allow for longer disease progression, showing increased maximum dilatation prior to reaching the rupture exit threshold. Results are shown for an axisymmetric insult distribution.

**Figure 7:**
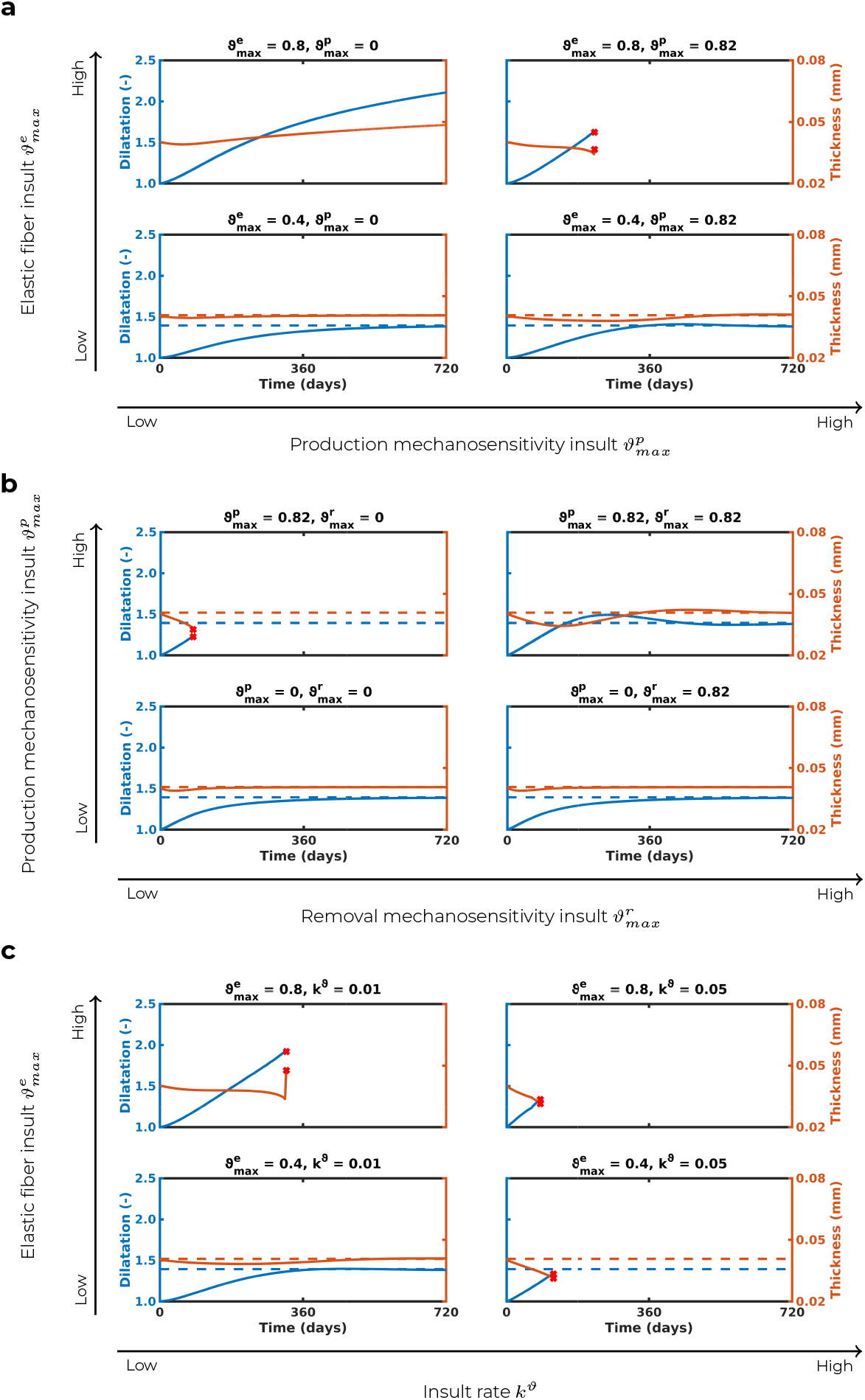
Effect of co-varying aneurysm insult parameters in vessels with a spatially asymmetrical insult profile. (a) Effect of co-varying elastic fiber insult and production mechanosensitivity insult. Removal mechanosensitivity insult is fixed at 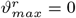 and insult rate is fixed at *k*^*ϑ*^ = 0.01. Increasing elastic fiber insult creates unstable growth while increasing production mechanosensitivity insult causes the rupture exit threshold to be reached. (b) Effect of co-varying production mechanosensitivity insult and removal mechanosensitivity insult. Elastic fiber insult is fixed at 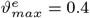 4 and insult rate is fixed at 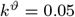. TAA stability is maintained with low 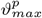 even with high values of 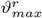 results in eventual . Conversely, high 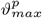 results in eventual loss of stability, although simultaneously increasing 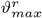 can allow stability to be maintained. (c) Effect of co-varying elastic fiber insult and insult rate. Production mechanosensitivity insult is fixed at 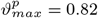 and removal mechanosensitivity insult is fixed at 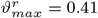. Increasing *k*^*ϑ*^ results in the exit rupture threshold being reached, even at moderate values of 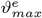 that would be expected to stabilize at low *k*^*ϑ*^. At severe 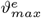 values, low values of *k*^*ϑ*^ allow further disease progression with increased maximum dilatation prior to reaching the simulation exit threshold.

**Figure 8:**
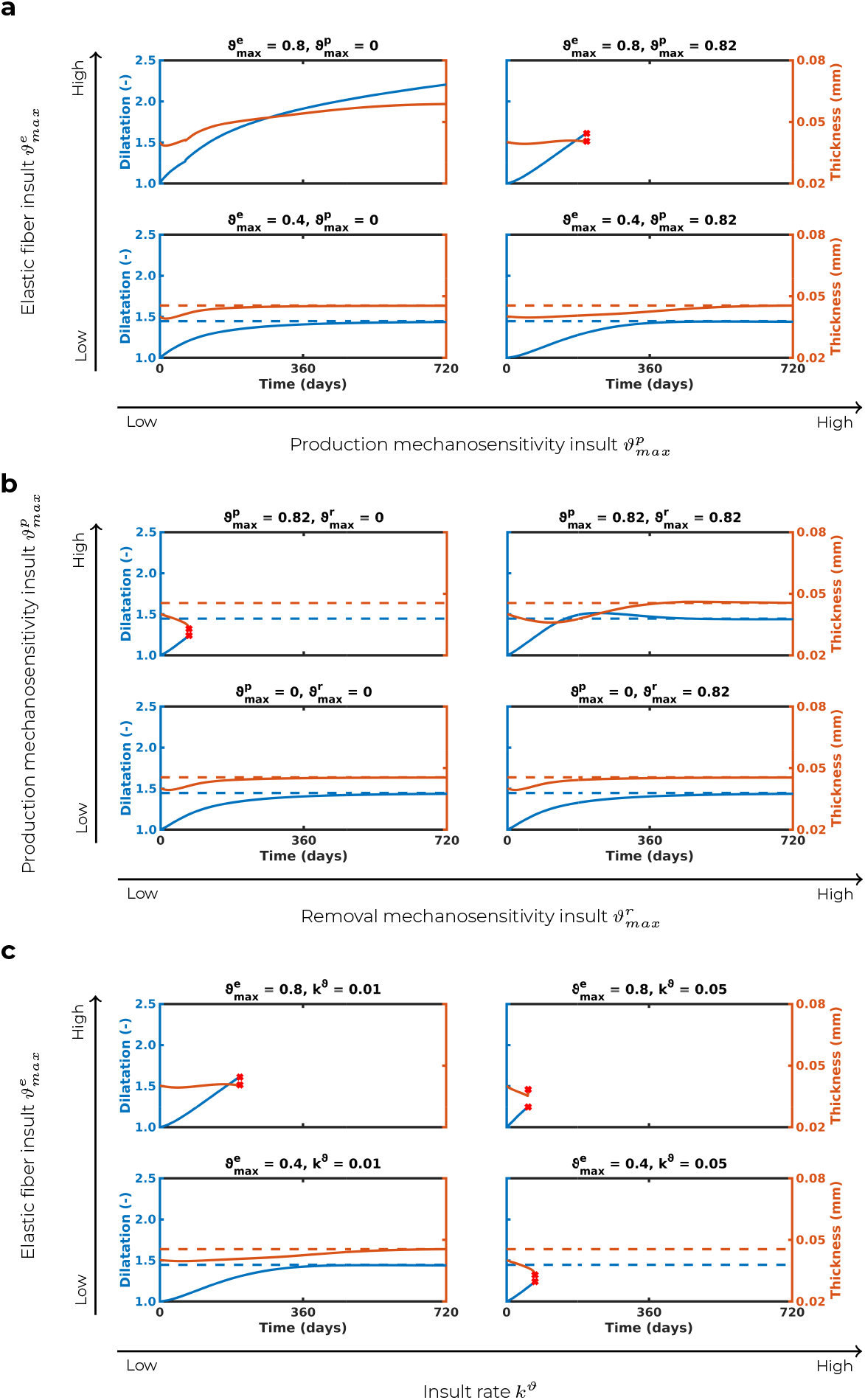
Effect of co-varying aneurysm insult parameters in vessels with a spatially random insult profile. (a) Effect of co-varying elastic fiber insult and production mechanosensitivity insult. Removal mechanosensitivity insult is fixed at 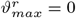 and insult rate is fixed at *k*^*ϑ*^ = 0.01. Increasing elastic fiber insult creates unstable growth while increasing production mechanosensitivity insult causes the rupture exit threshold to be reached. (b) Effect of co-varying production mechanosensitivity insult and removal mechanosensitivity insult. Elastic fiber insult is fixed at 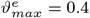 and insult rate is fixed at 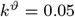. TAA stability is maintained with low 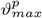 even with high values of 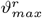. Conversely, high 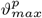 results in eventual loss of stability, although simultaneously increasing 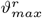 can allow stability to be maintained. (c) Effect of co-varying elastic fiber insult and insult rate. Production mechanosensitivity insult is fixed at 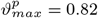 and removal mechanosensitivity insult is fixed at 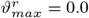. Increasing *k*^*ϑ*^ results in the exit rupture threshold being reached, even at moderate values of 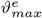 that would be expected to stabilize at low *k*^*ϑ*^. At severe 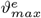 values, low values of *k*^*ϑ*^ allow further disease progression with increased maximum dilatation prior to reaching the simulation exit threshold.

### Elastic fiber insult increases low-risk TAA size, while production mechanosensitivity insult increases rupture risk

Dilatation increased with higher 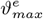 and 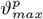 (Figure 4), with 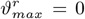 and *k*^*ϑ*^ = 0.010 held constant across these simulations. For cases where 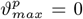, the time-resolved framework converged asymptotically toward the MBE predicted values for all metrics, supporting the validity of using the MBE for TAA evolutions with perfect mechanosensitivity (Figure 4a–c, top left). For moderate elastic fiber insult (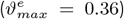), both formulations resulted in the vessel dilatation stabilizing at the threshold for aneurysmal classification (*>*1.5 times the original radius) with modest thickening (Figure 4a), as well as increased circumferential stiffness (Figure 4b), decreased strain energy, and well-regulated intramural stress (Figure 4c). Once the elastic fiber integrity insult surpassed 70%, stability of the TAA was lost in the time-resolved model, giving rise to an unbounded growth behavior, for which no corresponding solution existed in the MBE (Figure 4a–c, bottom left).

For cases where the production mechanosensitivity was severely compromised, the time-resolved model exited before the halfway point in the time-course, even at moderate elastic fiber insult values (Figure 4a–c, top right). This effect was exacerbated with increasing values of 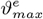, with the simulations reaching the rupture exit threshold within a quarter of the full time-course (Figure 4a–c, bottom right). Additionally, the computed metrics no longer approached those predicted by the MBE, frequently overestimating the stable value. Aneurysms with compromised production mechanosensitivity 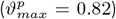 exhibited uncontrolled increase in circumferential stiffness and stress as compared to their perfect-mechanosensitivity counterparts. Additionally, in the case of moderate elastic fiber insult 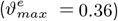, there was an increase in strain energy prior to exiting that differed strikingly from the expected MBE behavior (Figure 4c), though in severe elastic fiber insult case, strain energy decreased before exiting.

### High production and removal insult prolong unstable growth

To examine the relationship between production and removal mechanosensitivity, we fixed 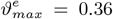 and *k*^*ϑ*^ = 0.050 while co-varying 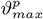 and 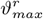 between 0 (perfect mechanosensitivity) and 0.82 (severe mechanosensitivity loss). For perfect mechanosensitivity, we obtained the expected convergence of the time-resolved solution to the MBE prediction. As long as production mechanosensitivity remained intact, the two frameworks continued to produce consistent predictions, even with severely compromised removal (Figure 5a–c, top rows). The time-resolved simulation showed high sensitivity to insults in production mechanosensitivity, in which any simulation with a high value of 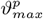 rapidly lost convergence.

Prescribing high insults to both production and removal allowed the simulation to progress further, slowing the course of disease progression and rupture, resulting in overall larger aneurysms with higher stresses and stiffness before reaching the rupture exit threshold (Figure 5a–c, bottom right). All cases of severe production mechanosensitivity insult resulted in the rupture exit threshold being reached in what was originally a small, stable TAA in the perfect mechanosensitivity and MBE models (Figure 5a–c, bottom rows). As before, we also observed uncontrolled increases in stiffness and stress exceeding the MBE predictions (Figure 5b, bottom right), as well as increases in strain energy along with poorly regulated intramural stress (Figure 5c, bottom right).

### Reduced insult rate can recover TAA stability

Finally, we investigated the relationship between the severity of elastic fiber insult and the insult rate by co-varying 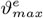 and *k*^*ϑ*^ while fixing the production and removal mechanosensitivity insults at 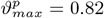 and 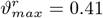, which are expected to result in unstable outcomes (Figure 3). Decreasing the rate at which elastic fiber and mechanosensitivity insults occurred (*k*^*ϑ*^ = 0.010) could stabilize the TAA, compared to the same simulations with rapidly evolving insults (Figure 5). Indeed, the moderately compromised evolution with a slow insult rate was shown to gradually converge to the MBE prediction (Figure 6a–c, top left). However, the time-resolved model continued to show that elastic fiber insult values beyond 70% could not reach a stable configuration even with a significantly lower insult rate (Figure 6, bottom rows), for which no MBE equivalent existed.

### Effects of insult profile asymmetry

Variations in insult profile spatial distribution all exhibited the same qualitative trends, with differences emerging in the insult magnitudes at which stability would be lost.

#### Asymmetric Insults

Asymmetric insult profiles are preferentially applied to one side of the aortic domain, as represented by Equation 11. This resulted in non-uniform dilatation that is representative of in vivo TAA cases which are often asymmetric. It has previously been shown that limiting the circumferential spread of insult enables higher maximum elastic fiber insults to be prescribed before stability is lost (Latorre and Humphrey, 2020b). Within the present framework, this allows an insult severity of 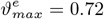 before the MBE formulation is unable to provide a stabilized solution. This higher threshold for instability was reflected in the outcomes of the time-resolved simulations. Unlike the symmetric insult profiles, a stable configuration was reached at high production mechanosensitivity insults with moderate elastic fiber insults in the asymmetric insult profiles (Figure 7a–c, top right). However, high elastic fiber insults displayed similar trends to those in symmetric insult profiles. High elastic fiber insult in the absence of a mechanosensitivity insult resulted in an unstable, growing aneurysm that did not reach the rupture threshold criteria during the simulation time-course. At high production mechanosensitivity insult, high elastic fiber insult reached the rupture threshold criteria approximately halfway through the simulation. Higher removal insult and lower insult rate prolonged the timecourse and final magnitude of unstable aneurysm growth prior to reaching the rupture threshold criteria (Figure 7). Like symmetrical aneurysm insults, stable outcomes agreed with MBE predictions in the asymmetrical cases.

#### Random Insults

Random insult profiles resulted in irregular, more natural TAA geometries. Aneurysms initiated with these insults tended to lose stability along a similar time-course as the idealized asymmetric cases. This similarity demonstrates that a lack of circumferential symmetry may result in similar stability behavior, even in more complex lesions (Figure 8). Random insult profiles with high elastin insult resulted in unstable growth. At high production mechanosensitivity insult, high elastic fiber insult resulted in the rupture threshold criteria being reached. High removal mechanosensitivity insult and low insult rate both resulted in a prolonged time-course and magnitude of aneurysm growth, resulting in overall larger dilatation prior to rupture. Stable outcomes in vessels with random insult profiles agreed with MBE predictions.

## DISCUSSION

Not all TAAs are the same. About 20% arise from predisposing pathogenic variants (e.g., *FBN1* variants that give rise to Marfan syndrome, MFS) or congenital defects (primarily bicuspid aortic valve, BAV), with the remaining *∼*80% arising sporadically, often in older individuals. A 2016 meta-analysis of growth rates of TAAs and associated risk factors assessed data from 11 publications (1,383 patients total) and found overall mean rates of lesion enlargement ranging from 0.2 to 4.2 mm/yr, with accelerated growth noted in larger lesions as well as in MFS and BAV patients (Oladokun et al., 2016). The authors concluded, nonetheless, that there was “a shortfall in the understanding of TAA expansion rates.” A more recent meta-analysis based on 55 publications further showed the importance of evaluating growth in different classes of lesions, with growth rates of 0.81 mm/yr in Loeys-Dietz syndrome, 0.45 mm/yr in MFS, 0.37 mm/yr in BAV patients, 0.33 mm/yr in sporadic TAAs, and 0.25 mm/yr in Turner syndrome (Henry et al., 2025). Clearly, there is a need to understand better the underlying mechanisms that drive such different growth rates in different patient populations. It is suggested herein that computational models can contribute in this regard.

Most prior computational studies of TAAs have focused on the hemodynamics (Caballero and Laín, 2013; Febina et al., 2018), calculation of wall stresses using finite element analyses (Martin et al., 2015; Wang et al., 2021), or modeling fluid-solid interactions (Mousavi et al., 2021; Pasta et al., 2013). These studies, and many others like them, have provided significant insights into the biomechanical complexities of TAAs but have not provided detailed information on mechanisms that drive progressive lesion enlargement versus arrest. One counterexample, however, is a finite element study wherein biaxial wall stresses were computed in TAAs for patients based on serial medical imaging over 1-to 6-year periods (Zamirpour et al., 2024). This study suggested that evolving values of wall stress are better predictors of all-cause mortality than evolving values of aortic diameter, but there were few details provided on the assumed material properties and their likely evolution. A more natural way to study progressive lesion enlargement is to use computational G&R models. Such data-informed models have been proposed recently for TAAs both for patients (Ghavamian et al., 2020; Mousavi et al., 2019) and for mice (Bazzi et al., 2025; Li et al., 2023), though without detailed parameter sensitivity studies that could be useful in understanding potential effects of underlying contributors to disease (Baek et al., 2005; Brandstaeter et al., 2021; Valentin et al., 2011).

In this paper, we extended prior G&R simulations that focused on potential mechanobiological drivers of TAA (Latorre and Humphrey, 2020b) through inclusion of rate-based parameters within constrained mixture models of vascular growth and remodeling. We focusced on the striking influence of mechanosensitivity factors on TAA phenotype, stability, and outcome. We also showed that the more computationally efficient MBE formulation indeed offers sound predictions of time-resolved outcomes when employed in scenarios where a mild insult severity and gradual rate, coupled with robust mechanosensitivity, may eventually stabilize the growth. This is the first time such a comparison has been made across three-dimensional finite element models; it demonstrates that the MBE framework identifies unique stable configurations, as long as such states exist. Given that the MBE framework is significantly less computationally expensive than time-resolved simulations, this finding supports use of the MBE framework for studies when identifying stable configurations under robust mechanosensing and mechanoregulation is expected. However, it also demonstrates the critical importance of considering rate-dependent parameters not able to be captured by the MBE framework, as these contribute significantly to simulated aneurysm outcomes.

Our idealized axisymmetric and asymmetric insult profiles illustrated general trends in how structural (*ϑ*^*e*^), mechanobiological (*ϑ*^*p*^ and *ϑ*^*r*^), and rate-based (*k*^*ϑ*^) parameters impacted aneurysm growth and stability. Not surprisingly, greater elastic fiber insult resulted in larger aneurysms. At very high elastic fiber insult values, there was no achievable stable configuration, though predicted by the MBE framework, and the aneurysm continued to grow past the simulation endpoint. Despite the unbounded growth due to large elastic fiber insults, an additional insult to production mechanosensitivity was still required to reach a rupture threshold in our simulation time-frame. At large production mechanosensitivity insults, larger dilatations, stresses, and stiffnesses were produced for the same magnitudes of elastic fiber insult compared to perfect mechanosensitivity counterparts. Even moderate elastic fiber insults that would otherwise result in stable, non-aneurysmal outcomes could be destabilized by a severe production mechanosensitivity insult to achieve rupture exit threshold outcomes. By simultaneously varying the removal mechanosensitivity insult, this behavior could be tuned to achieve rupture at either a small diameter or a large diameter, demonstrating how variations in underlying cellular processes can result in different rupture conditions. This observation held in asymmetric and random insult patterns, although the reduced circumferential spread of the lesion tended to allow more severe insults and longer time-courses to be simulated before reaching the rupture exit threshold. The insight gained from the simplest axisymmetric profile could offer a path forward for more computationally tractable parametric studies of aneurysm G&R without as great of a need for high-fidelity geometric representations.

These simulations also allow quantification of the biomechanical behavior of aneurysms that may not be feasible to measure in vivo and may identify potential targets for further experimental investigation. For example, experimental studies of aneurysms in *Fbn1*^*C1041G/+*^ and *Fbn1*^*mgR/mgR*^ mice showed higher circumferential stiffness and lower strain energy in TAAs (Bellini et al., 2016; Cavinato et al., 2021). In our computational study, stable and growing dilatations that did not rupture agreed with this observation. However, our simulations also explored a class of lesions with moderate elastic fiber insult and high production mechanosensitivity insult that exhibited increased stiffness, stress, and strain energy before eventual rupture. Although ruptured aortas cannot be biomechanically evaluated experimentally, this modeling framework offers some predictions of the pre-rupture biomechanical phenotype, though this would be difficult to validate. Nevertheless, this may suggest that certain trends in decreasing strain energy could be indicative of underlying mechanisms of aneurysm progression and risk of rupture.

Establishing a three-dimensional framework capable of generating time-resolved simulations of (i) stable aortic dilatation, (ii) progressive dilatations, (iii) small-diameter rupture, and (iv) large-diameter rupture is of significant clinical value. These categories parallel clinical decision-making, where distinguishing between stable and unstable cases — even among lesions with similar intermediate geometries — is central to risk assessment and treatment planning. Demonstrating that these outcomes can be reliably produced in a time-resolved framework for both idealized and complex insult profiles establishes a robust foundation for a variety of in silico studies. Looking forward, whereas regression models may prove increasingly useful in predicting which individuals are at risk of developing TAA (Mori et al., 2021), or perhaps even which patients are susceptible to unstable lesion growth, the ability to generate large amounts of physiologically-relevant synthetic data is particularly relevant to large data models in the clinical setting for diagnosis and treatment planning. For example, machine learning models often rely on large high-quality datasets on which to train. Such clinical datasets are rarely available, but a robust simulation platform can help overcome this limitation by generating synthetic datasets that mimic in vivo behavior. This approach has been used in transfer learning, where simulated data augments limited clinical data to substantially improve neural operator performance (Goswami et al., 2022).

Notwithstanding new insight gleaned from the simulations reported herein, many additional considerations remain. We evaluated time-course changes in geometric and mechanical metrics of interest in the region of maximum insult to evaluate possible stabilization of lesion growth, but we did not perform a formal mechanobiological stability analysis (Cyron and Humphrey, 2014; Latorre and Humphrey, 2019); such analysis of TAA growth could provide additional insight (cf. Cyron et al. (2014)). We also did not consider initial regional variations in material properties, though such differences exist in both humans (Davis et al., 2016) and mice (Bersi et al., 2019). There is a pressing need for information on regional properties and how they may evolve with lesion growth (cf. Cavinato et al. (2021)), and the present three-dimensional finite element platform is well suited for such investigations. Additionally, we focused solely on phenomenological constrained mixture models of vascular G&R. Such models have proven effective in describing and predicting tissue-level changes in diverse cases of vascular disease (Humphrey, 2021); nevertheless, there are considerable advantages to coupling tissue-level G&R models with cellular signaling models (Irons et al., 2021) to consider both fundamental mechanisms and potential therapeutic targets.

## CONCLUSION

Computational simulations have the potential to aid clinical decision-making for TAAs by revealing how diverse biomechanical factors influence stabilization, growth, and rupture. By incorporating additional rate-based parameter insults into the constrained mixture theory of G&R, we developed a computational framework capable of simulating clinically relevant outcomes of TAAs. This approach enables us to identify biomechanical trends that precede each outcome type and examine how combinations of insult types produce distinct phenotypes. Furthermore, we directly compared outcomes from time-resolved constrained mixture simulations to MBE simulations, demonstrating that the MBE framework robustly predicts stabilized configurations when such outcomes exist. These findings motivate use of the MBE approach– which is an order of magnitude more computationally efficient– for identifying and simulating stable outcomes, while reserving the time-resolved framework for cases when the time-course evolution or instability of vessels must be captured. Altogether, this work establishes a foundation for clinically relevant in silico studies with translational potential.

## ACKNOWLEDGMENTS

This work was supported in part by grants from the NIH (R01 HL168473, R01 HL169147, and P01 HL169168). This work used Expanse at the San Diego Supercomputer Center through allocation MDE230014 from the Advanced Cyberinfrastructure Coordination Ecosystem: Services & Support (ACCESS) program, which is supported by U.S. National Science Foundation grants 2138259, 2138286, 2138307, 2137603, and 2138296 (Boerner et al., 2023).

