## Supplementary Materials for "Mechanisms Driving Thoracic Aortic Aneurysm Stability"

**This PDF file includes:**

Figs. S1 to S4

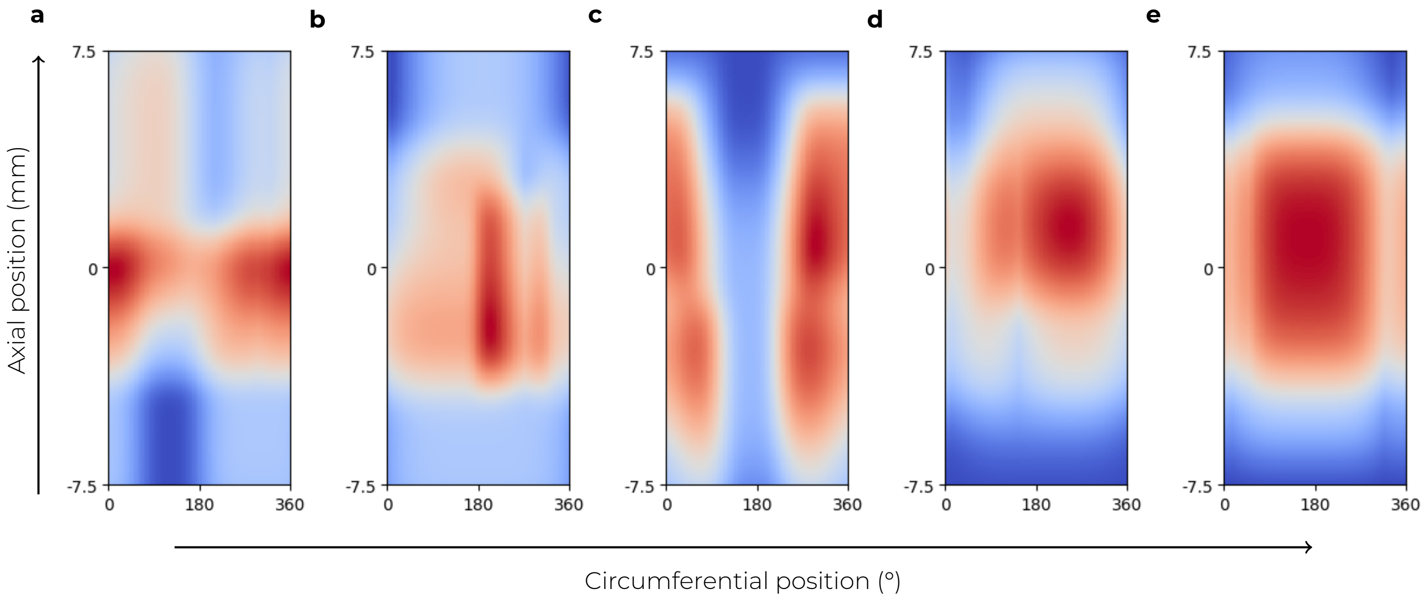


**Figure S1: Examples of randomly generated insult fields.** Spatial insult profiles generated using the process described in Methods. Fields are periodic in the circumferential direction. Field (b) was used in this paper for the random insult spatial profile.

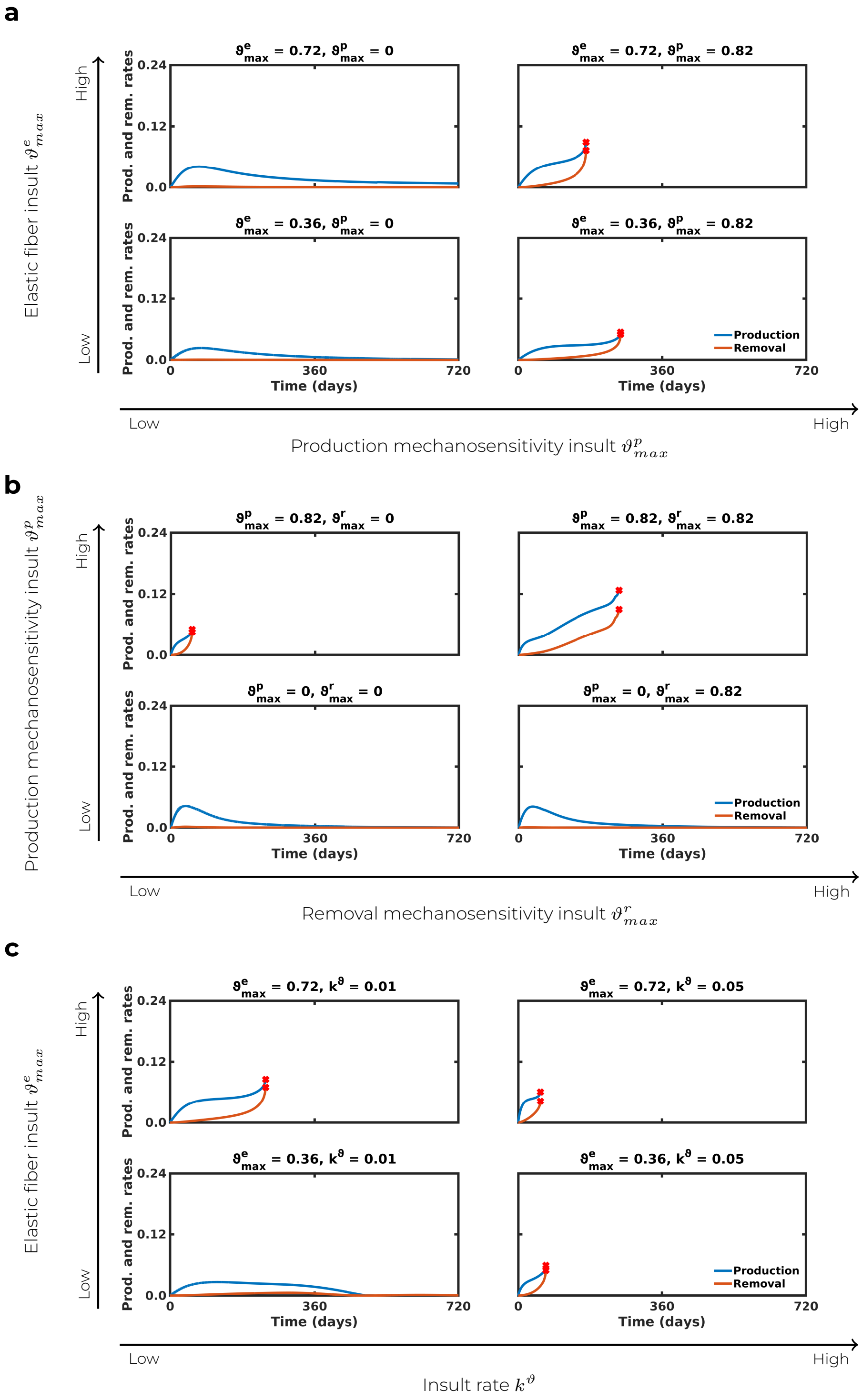


**Figure S2: Removal and production stimuli function for co-varying insults in a vessel with a spatially axisymmetrical insult profile.** Production $((1-\vartheta^p(\tau))\,K^{\alpha}_{\sigma}\Delta\sigma(\tau)-(1-\vartheta^{\tau}(\tau))\,K^{\alpha}_{\tau_w}\Delta\tau_w(\tau))$ and removal $((1-\vartheta^r(\tau))\,\omega^{\alpha}_{\sigma}\Delta\sigma^2$(t)) rates associated with (a) Figure 4, (b) Figure 5, and (c) Figure 6. Simulation exit threshold is generally met when removal rate overtakes, or begins to overtake, the production rate.

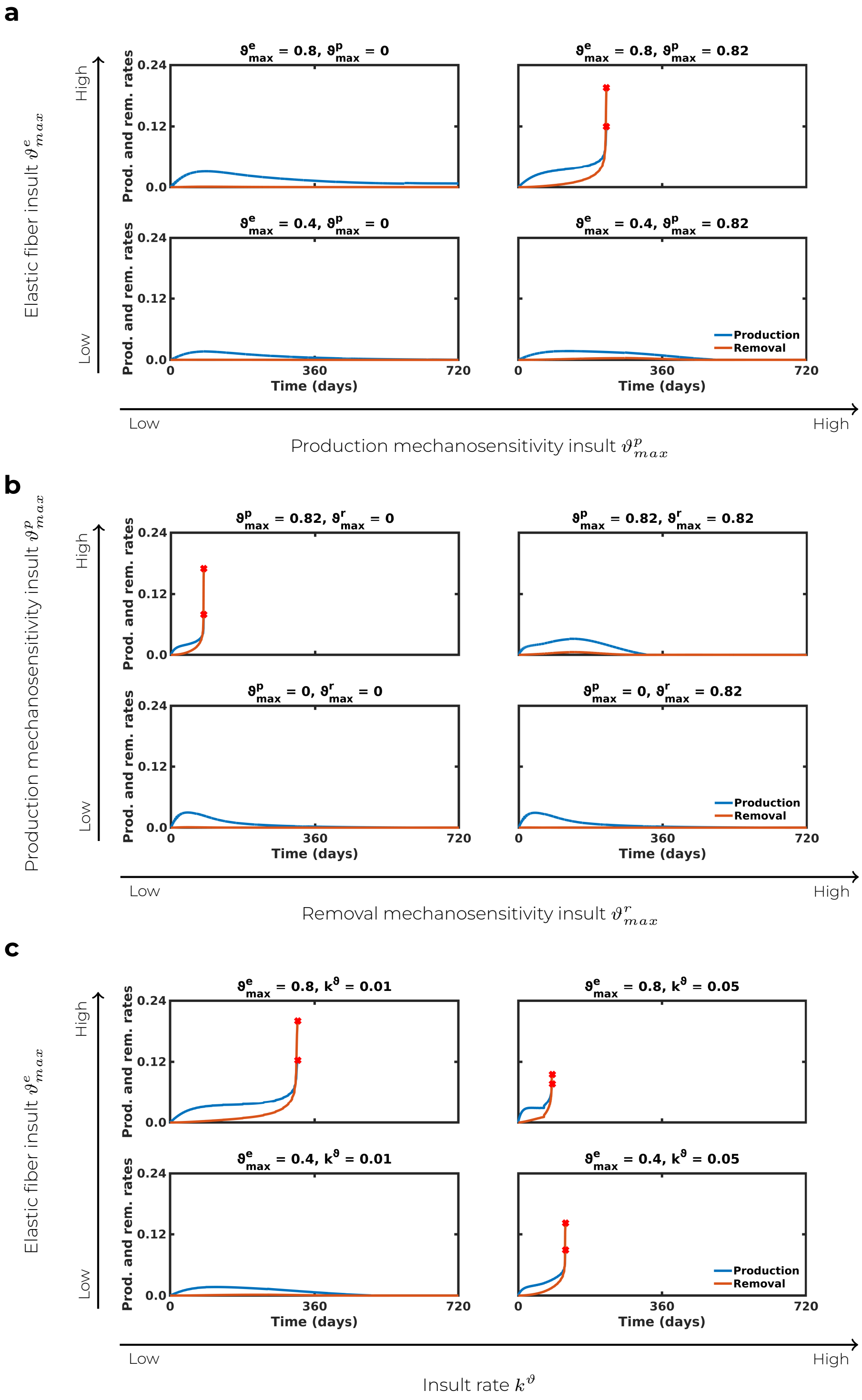


**Figure S3: Removal and production stimuli function for co-varying insults in a vessel with a spatially asymmetrical insult profile.** Production $((1-\vartheta^p(\tau))\,K^\alpha_\sigma\Delta\sigma(\tau)-(1-\vartheta^\tau(\tau))\,K^\alpha_{\tau_w}\Delta\tau_w(\tau))$ and removal $((1-\vartheta^r(\tau))\,\omega^\alpha_\sigma\Delta\sigma^2$(t)) rates associated with (a) Figure 7, (b) Figure 8, and (c) Figure 9. Simulation exit threshold is generally met when removal rate overtakes the production rate.

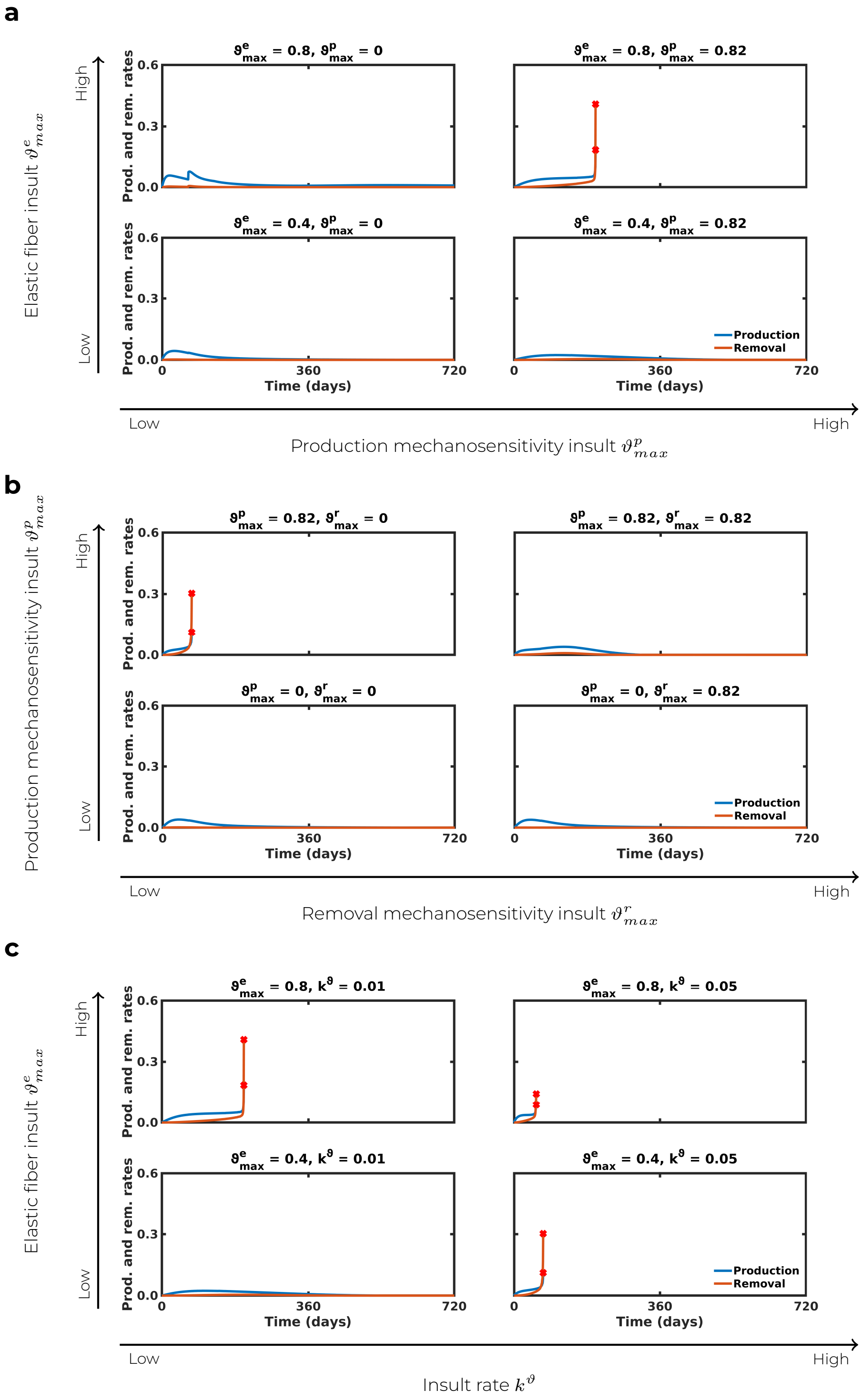


**Figure S4: Removal and production stimuli function for co-varying insults in a vessel with a spatially random insult profile.** Production ($(1-\vartheta^p(\tau))\,K^\alpha_\sigma \Delta\sigma(\tau) - (1-\vartheta^\tau(\tau))\,K^\alpha_{\tau_w}\Delta\tau_w(\tau)$) and removal ($(1-\vartheta^r(\tau))\,\omega^\alpha_\sigma \Delta\sigma^2$(t)) rates associated with (a) Figure 10, (b) Figure 11, and (c) Figure 12. Simulation exit threshold is generally met when removal rate overtakes the production rate.
